# The Xenopus Phenotype Ontology: bridging model organism phenotype data to human health and development

**DOI:** 10.1101/2021.11.12.467727

**Authors:** Malcolm E. Fisher, Erik Segerdell, Nicolas Matentzoglu, Mardi J. Nenni, Joshua D. Fortriede, Stanley Chu, Troy J. Pells, Praneet Chaturvedi, Christina James-Zorn, Nivitha Sundararaj, Vaneet S. Lotay, Virgilio Ponferrada, Dong Zhuo Wang, Eugene Kim, Sergei Agalakov, Bradley I. Arshinoff, Kamran Karimi, Peter D. Vize, Aaron M. Zorn

**Affiliations:** Division of Developmental Biology, Cincinnati Children’s Hospital Medical Center, Cincinnati, Ohio, USA; Monarch Initiative; Semanticly Ltd, London, UK; European Bioinformatics Institute (EMBL-EBI), UK; Department of Biological Science, University of Calgary, Calgary, Alberta, Canada

**Keywords:** Xenopus, Phenotypes, Ontology, disease models

## Abstract

**Background:** Ontologies of precisely defined, controlled vocabularies are essential to curate the results of biological experiments such that the data are machine searchable, can be computationally analyzed, and are interoperable across the biomedical research continuum. There is also an increasing need for methods to interrelate phenotypic data easily and accurately from experiments in animal models with human development and disease.

**Results:** Here we present the *Xenopus* Phenotype Ontology (XPO) to annotate phenotypic data from experiments in *Xenopus*, one of the major vertebrate model organisms used to study gene function in development and disease. The XPO implements design patterns from the Unified Phenotype Ontology (uPheno), and the principles outlined by the Open Biological and Biomedical Ontologies (OBO Foundry) to maximize interoperability with other species and facilitate ongoing ontology management. Constructed in Web Ontology Language (OWL) the XPO combines the existing uPheno library of ontology design patterns with additional terms from the *Xenopus* Anatomy Ontology (XAO), the Phenotype and Trait Ontology (PATO) and the Gene Ontology (GO). The integration of these different ontologies into the XPO enables rich phenotypic curation, whilst the uPheno bridging axioms allows phenotypic data from *Xenopus* experiments to be related to phenotype data from other model organisms and human disease. Moreover, the simple post-composed uPheno design patterns facilitate ongoing XPO development as the generation of new terms and classes of terms can be substantially automated.

**Conclusions:** The XPO serves as an example of current best practices to help overcome many of the inherent challenges in harmonizing phenotype data between different species. The XPO currently consists of approximately 22,000 terms and is being used to curate phenotypes by Xenbase, the *Xenopus* Model Organism Knowledgebase, forming a standardized corpus of genotype-phenotype data that can be directly related to other uPheno compliant resources.

## Background

Laboratory organisms, such as frogs, mice, fish, fruit flies and worms are essential to investigate conserved gene function and model human development, homeostasis and disease. In order for the experimental results in hundreds of thousands of published animal model papers to be computer readable, amenable to computational analysis and interrelated across species, the data must be curated using ontologies of controlled vocabulary describing genes, gene products, molecular and biological processes, anatomy, and phenotypes. Ontologies codify semantic relationships between biological concepts and are essential because natural language descriptions in publications are too variable, organism specific and cumbersome to be machine processed efficiently [1].

When biomedical ontologies initially began to proliferate the Open Biomedical Ontologies (OBO) consortium established a set of principles for ontology development to improve accessibility, specificity, and interoperability [2]. Curating with ontologies allows data from two papers using different phrases or synonyms to describe the same structure or phenotype (e.g. enlarged heart versus cardiac hypertrophy) to be annotated using the same ontology term and ID number. For a computer, the two experiments both use the same ID number and therefore contain data on the same anatomical structure. Ontologies also define the relationships between terms - for example the ‘heart’ is a part of the ‘cardiovascular system’ which also has its own unique ID number, thus papers on different parts of the cardiovascular system can be mapped to each other by a computer using this ‘part_of’ relationship ID. Ontologies can store many such relationships along with synonyms and cross-reference IDs to other ontologies, such as cellular components in an anatomy ontology being cross referenced to the equivalent Cell Component term in the Gene Ontology (GO) (RRID:SCR_002811) [3]. By curating data from thousands of publications with interconnected ontologies it is possible to build a web of knowledge that can be subjected to statistical and computational analyses. In the context of disease modeling, ontologies make it possible, in principle, to connect genotype and phenotype data from experiments in animal models with an understanding of pathological mechanisms associated with an orthologous human condition. To realize this potential, ontologies used by different biomedical research communities need to be interoperable and grounded in a common syntax.

Ontology based phenotype curation has traditionally taken one of two forms, either post-composed or pre-composed. In post-composed approaches two or more ontologies are used in combination, typically, an anatomical ontology term is used to define the entity (E) and this is combined with a quality ontology term (Q) to generate the entity-quality (EQ) phenotype description. For example, the entity may be ‘heart’ from an Anatomy Ontology and the quality may be ‘decreased size’ from the Phenotype and Trait Ontology (PATO) (RRID:SCR_004782) [4], these would be combined to generate the EQ statement: ‘heart, decreased size’. The entity component can be complex with multiple independent entity terms joined by Relationship Ontology [5] terms, such as ‘has_quality’ or ‘part_of‘, to give great flexibility in description. On the other hand, pre-composed approaches use a predefined phenotype ontology, where a single ontology term using a controlled syntax already exists, such as ‘decreased size of the heart’ (XPO:0103343). While post-composed annotation allows for richer detailed descriptions, it has the drawback that different curators may select different combinations of terms to describe the same phenotype thus increasing variability. A second drawback is that the different component ontologies often diverge, through the natural course of different groups working independently on ontology development, making synchronization a challenge and requiring frequent re-annotation to already curated data. This variability and lack of synchrony has made it difficult to make post-composed based phenotype assertions interoperable between different model organisms and humans. While variability is less of a problem with the pre-composed approach, one drawback is the scale of pre-composed ontologies – groups of terms must be in place for every phenotype and anatomical structure: a ‘small heart’, ‘small pancreas’, ‘small limb’, ‘small head’ etc. This makes the ontology very large as when almost every anatomy ontology term gives rise to multiple phenotype terms the phenotype ontology must be several times larger than the associated anatomy ontology. This large size is not a major issue as it can be tackled programmatically, both for generation and management. A final challenge shared by both pre- and post-composed approaches is that anatomical structure, and terms commonly used to describe them in the literature, can be very species-specific.

The fact that different research communities use distinct approaches and different ontologies to curate phenotypes has been a challenge for cross-species comparisons. For example, human clinical disease phenotyping is mostly done with the pre-composed Human Phenotype ontology (HPO) (RRID:SCR_006016)[6, 7], the Mammalian Phenotype ontology (MP) (RRID:SCR_004855) [8, 9]. The Zebrafish Information Network (ZFIN) (RRID:SCR_002560) [10] uses a post-composed EQ approach combining the Zebrafish Anatomical Ontology (ZFA) (RRID:SCR_005887) [11], GO and PATO[4]. An approach to harmonize different phenotype ontologies and enable cross-species comparisons was recently established by the Monarch Initiative; a multi-species bioinformatic resource aggregating genotype and phenotype data from multiple model organisms to inform the genetic basis of human disease [12, 13]. The Monarch consortium and their collaborators, which include most of the major model organism knowledgebases, implemented the Unified Phenotype Ontology (uPheno) that uses ‘bridging axioms’ to equate terms from different species-specific ontologies. Monarch produces a knowledge graph based representation using data and ontologies loaded into a SciGraph database [13] where entities (from different ontologies) are represented by nodes connected by edges representing distinct relationships. uPheno allows connectivity and equivalences between the phenotype ontologies of multiple species. An important component of the uPheno plan was a community wide effort to reconcile and align different ontologies using a standard pre-composed template to maximize interoperability [14].

Leveraging these recent advances, we set out to build a uPheno-compliant *Xenopus* phenotype ontology (XPO) that Xenbase, the *Xenopus* model organism knowledgebase (RRID:SCR_003280)[15, 16], could use to curate phenotype data from *Xenopus* experiments with maximum interoperability to humans and other model organisms. The frog, *Xenopus*, is one of the leading vertebrate model systems and has been a major contributor to understanding fundamental biological processes such as cell division, cell differentiation, morphogenesis, organogenesis and neurobiology. It is also increasingly used to model human disease, particularly congenital conditions [17]. As a tetrapod, *Xenopus* occupies a key evolutionary niche between fish and mammals. The large, abundant, externally developing *Xenopus* embryos have several unique features that lend themselves to functional genomics and disease modeling including; CRISPR gene editing, antisense morpholino knockdown, transgenics, experimental embryology, and live cell imaging. We estimate that there are ~4,000 publications with phenotype data from *Xenopus* experiments in Xenbase with more papers published every month, but up until recently there were no largescale efforts to curate this data and thus it was largely inaccessible to the wider biomedical community. Below we describe the development and implementation of a fully uPheno compliant XPO, which will facilitate access to *Xenopus* phenotypic data for the biomedical community.

## Results

### XPO design strategy

We wanted to design the XPO to capture the breadth of experimental phenotypes in the *Xenopus* literature which range from disrupted gastrulation to congenital malformations and limb regeneration in adult frogs. To assess this phenotypic range a team of four expert Xenbase curators annotated phenotypes from 200 *Xenopus* papers with a free form Entity-Quality (EQ) syntax in the Phenote software package [1] using the *Xenopus* Anatomy Ontology (XAO) (RRID:SCR_004337) [18], PATO [4], GO[3], Basic Formal Ontology (BFO)[19] and the Relations Ontology (RO)[5]. This initial set of 1078 EQ phenotype statements served as a seed to develop the XPO. From the EQ definitions we extracted combinations of high level XAO, GO and PATO terms that we wanted to incorporate into the XPO such as “anatomical structure”, “cell part”, “morphology”, “localised” and “anatomical space” and mapped these to existing uPheno patterns. Figure 1 shows an example of how an EQ curation, for an image from Gouignard et al. 2016[20] summarized as ‘decreased size of the eye’, was used to construct a generalized design pattern of ‘decreased size of anatomical structure’, which was then applied to generate new ‘decreased size’ XPO terms for each appropriate anatomical entity in the XAO. This process, identified 13 frequently used patterns (Supplementary Table 1) that were then submitted as new class requests to the ongoing Phenotype Ontologies Reconciliation Effort (PORE) [21]. For example, several patterns were developed relating to cilium motility in various ciliated tissues. Once these new patterns were validated and added to uPheno by the ontology development team, we implemented them in the XPO. In this way Xenbase curators contributed to the definition of phenotype patterns that are now reused across many other domains. This reiterative approach ensures that the XPO remains in synchrony with uPheno.

**Figure 1:**
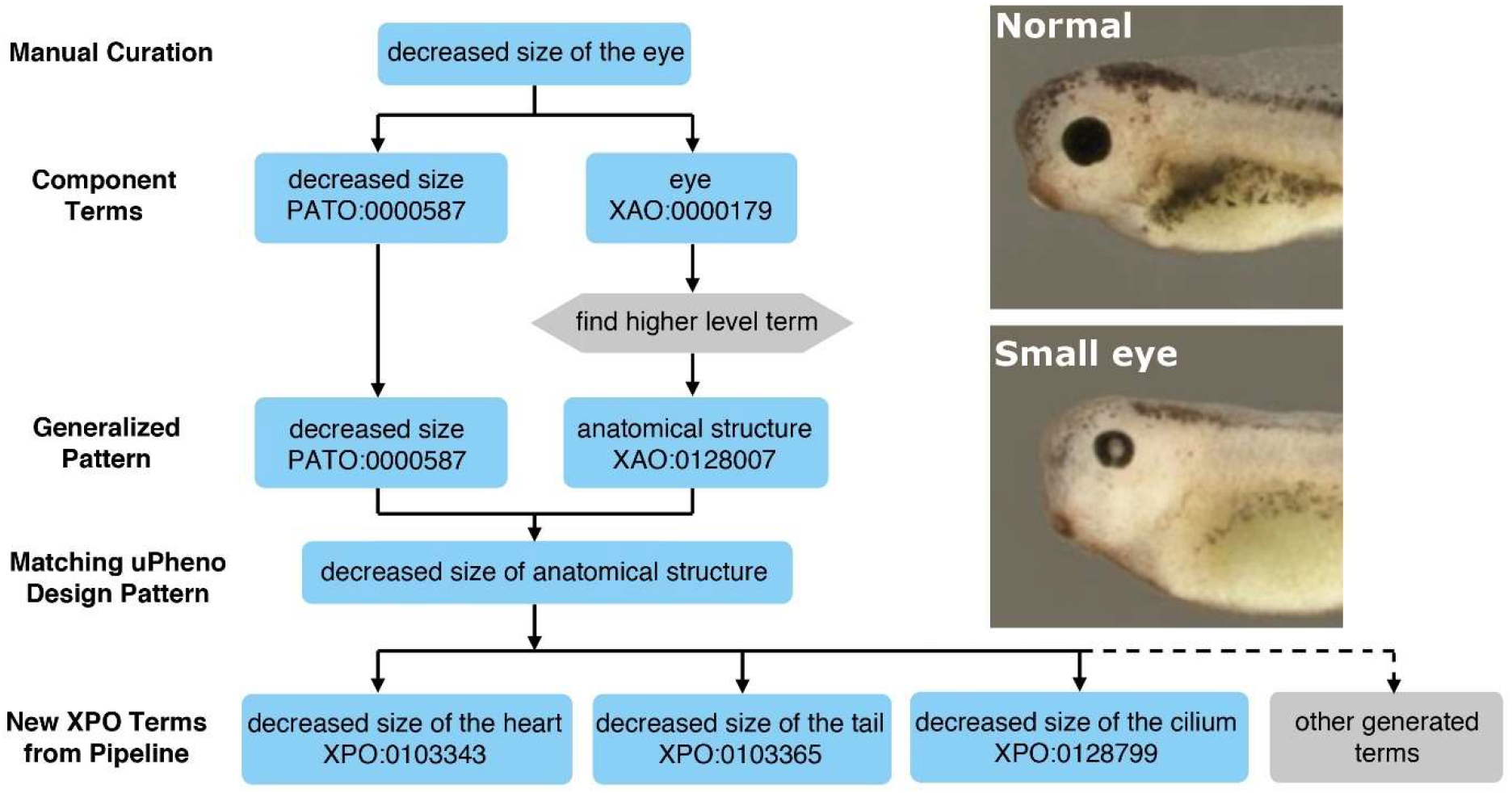
Workflow diagram for generating and applying a design pattern. Specific manually curated terms are decomposed and used to generate generalized patterns which are matched with existing uPheno design patterns or used for requesting new design patterns. The design patterns are then applied to tabulated sets of XAO, PATO or GO terms to generate new classes of XPO terms.

### Generating XPO terms using uPheno design patterns

To build a uPheno compliant XPO we used standard tools such as the Ontology Development Kit (ODK) [22] and OBO Tool (ROBOT) [23] to generate an ontology in the W3C Web Ontology Language (OWL) format [24], a semantic language designed for complex knowledge and relationships. By using uPheno design patterns as templates we were able to efficiently construct a pre-composed phenotype ontology incorporating terms from existing ontologies, such as the XAO, PATO, and molecular functions and biological processes from GO. uPheno design patterns prescribe a statement syntax which takes variables from a tab separated value (TSV) file containing a table of component terms to produce multiple appropriately formatted terms. Figure 2 shows a conceptual diagram of the pipeline including a partial example of a design pattern YAML file [25]. The pattern pipeline fills in new Internationalized Resource Identifiers (IRIs) in all TSV files corresponding to the selected terms. A second major step in the workflow, an automated release preparation pipeline, checks for updates to uPheno patterns and ontology, makes subclass assertions and generates uPheno-compliant logical definitions, flags errors such as duplicate equivalent classes and term labels, and builds the OWL files that comprise a new XPO release. Before making the release official, curators can inspect the ontology in an editor such as Protege to ensure that changes appear as expected. Curators may occasionally need to edit ontology metadata such as the ontology description by opening an “editor’s” OWL file; otherwise, routine class requests and updates are handled exclusively in the TSV tables and configuration file.

**Figure 2:**
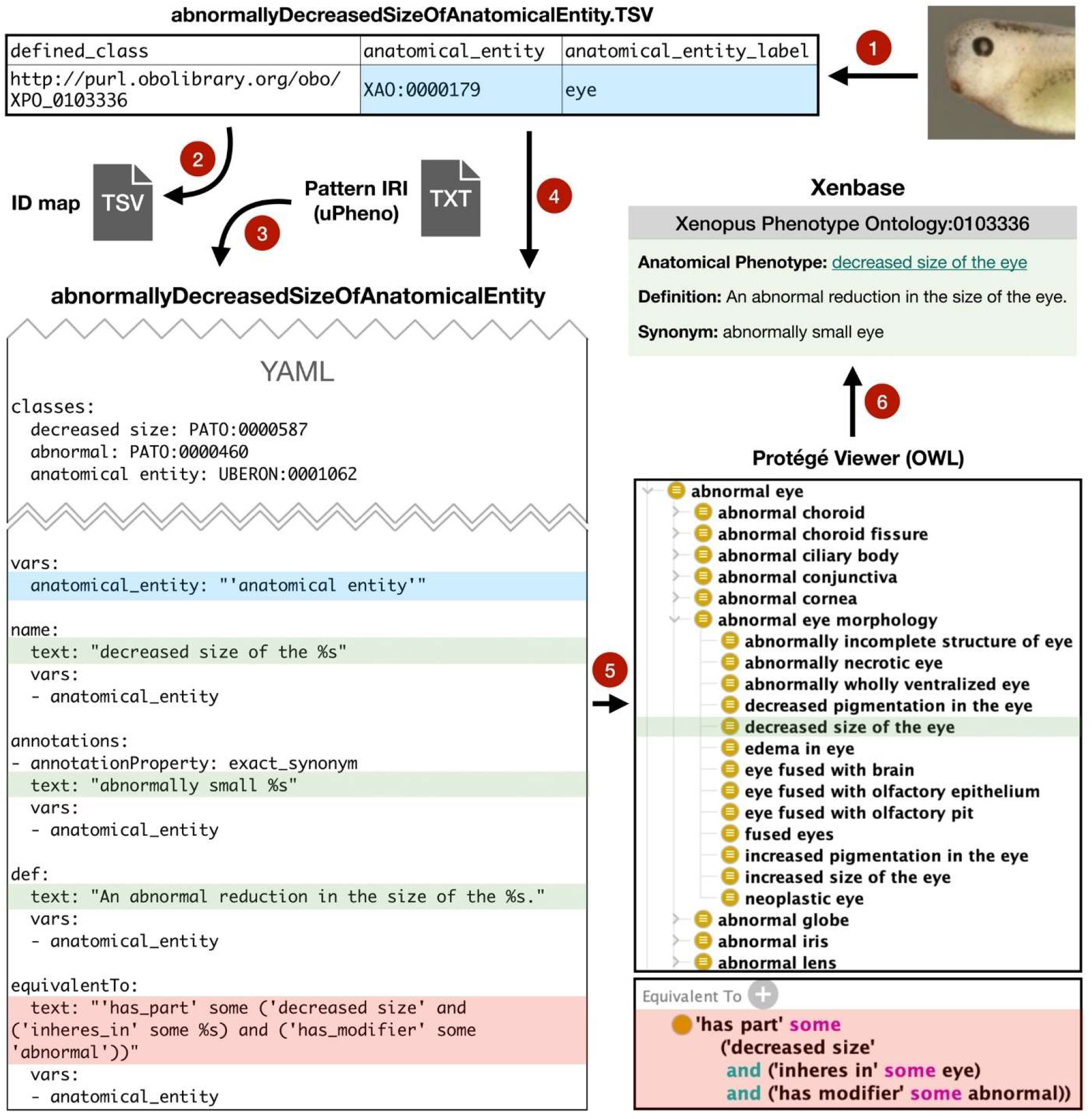
The building of an XPO term. XAO terms for phenotypes are selected (1) and entered in TSV files matched to specific design patterns (2). A partial uPheno design pattern YAML file example (3) shows the description of the pattern ‘Abnormally decreased size of anatomical entity’, and templates for generating names, synonyms, definitions, and equivalent classes for the pattern. The ‘%’ characters are substituted with the appropriate terms, in this case ‘anatomical entity’, from the TSV source tables during the ontology building process (4). The pipeline builds the new term from the specified component term and pattern and integrates it into the ontology (5), this shows the built equivalent classes with the XAO term variable filled. Once the new XPO build is complete with the new term it is made available on Xenbase (6).

The initial TSV lists for each pattern were generated based on the higher-level ontological classes from our previous review phase by identifying appropriate terms and selecting all their children with specific relationships. To enable more precise control over these automatically generated classes and prevent creation of phenotypes that make little or no biological sense some subclass terms and their children were excluded, for instance XAO terms that were children of ‘anatomical space’ were excluded from lists for the generic pattern ‘biological process in location’ for where the processes were the GO terms ‘cell population proliferation’ or ‘apoptotic process’, as these cellular processes do not occur in acellular anatomical spaces. Only certain relationships, such as ‘is_a’ and ‘part_of’ but not ‘develops_from’, are used when traversing from higher level to lower level XAO terms when selecting terms to be used in the XPO. ‘Develops from’ is not used as it would lead to the propagation of phenotypes in developed tissues to their precursor structures, which is not a given. While all ‘abnormal eyes’ are part of an ‘abnormal visual system’ an ‘abnormal eye’ does not necessarily imply that it developed from an ‘abnormal optic vesicle’. This leads to the structure of the XPO reflecting but not duplicating the XAO (Fig. 3).

**Figure 3:**
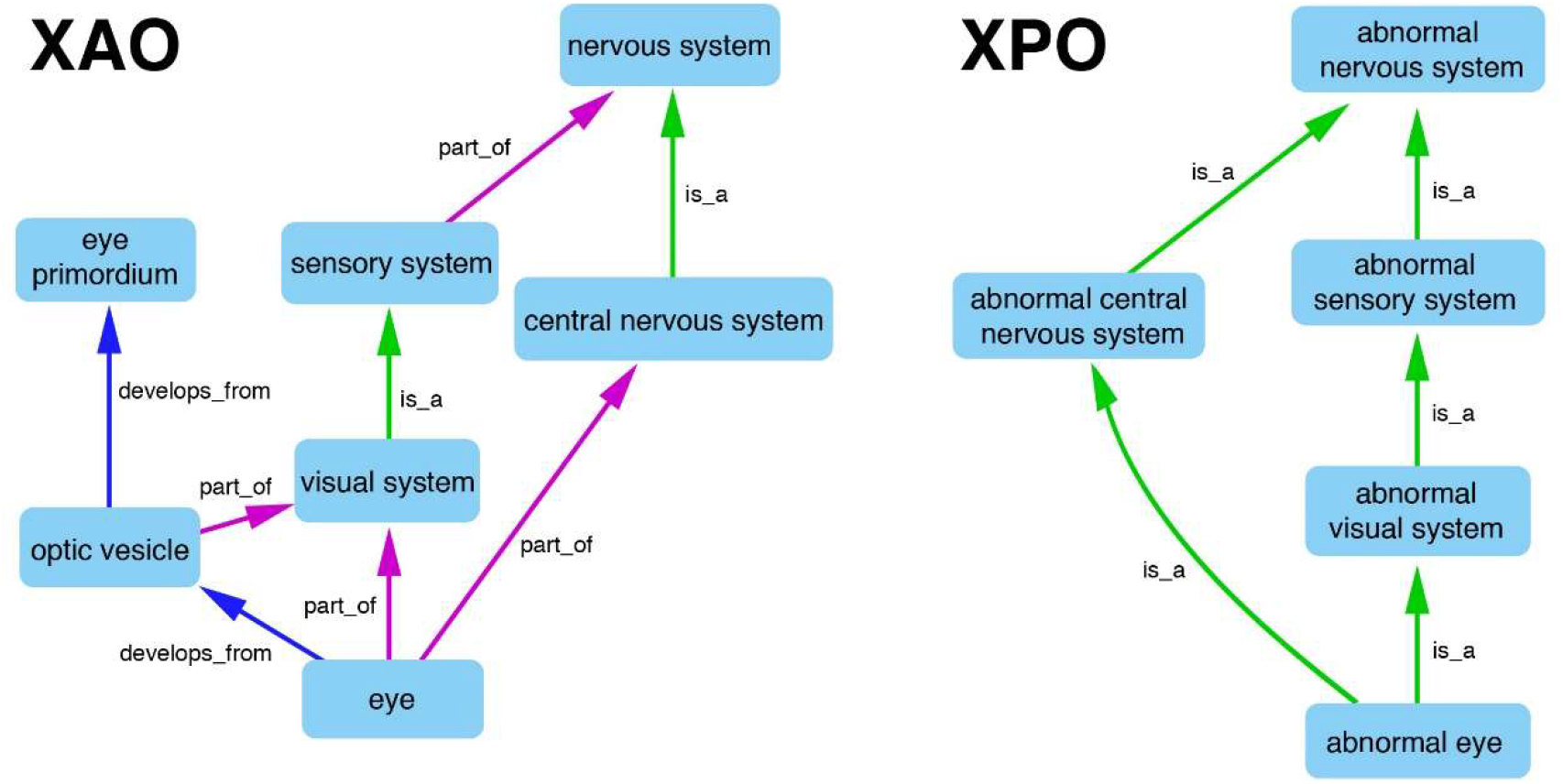
Comparison of graph visualizations of sections of the *Xenopus* anatomy ontology (XAO) and *Xenopus* phenotype ontology (XPO). The XPO structure reflects but does not reduplicate that of the XAO as only ‘part-of’ and ‘is_a’ XAO relationships are incorporated and the XPO uses only ‘is_a’ relationships. Consequently, relationships such as ‘develops_from’ are lost. Relationship edges are directed as indicated by arrows.

The XPO is the only phenotype ontology to date whose classification is purely driven by logical definitions and fully uPheno compliant. Subclass assertions, the defining of hierarchical parent child relationships between terms, a process that is known to be error-prone and incomplete, do not need to be made manually. This significantly reduces maintenance effort and increases interoperability of the XPO. We extended this novel approach to automatically construct phenotype terms from other ontologies, most importantly the XAO but also GO and NBO. For example, instead of having to curate a new phenotype term for “abnormal structure-*X*” whenever a new anatomy term is added to the XAO, the XPO pipeline automatically scans the XAO for new terms which are then used to automatically generate new XPO terms according to the specific predesignated design patterns (Table 1), these 14 standard patterns account for ~96% of terms in the XPO. This significantly reduces the effort in maintaining an up-to-date phenotype ontology and maintains synchrony with the anatomy ontology. We estimate that these patterns are likely to be used for comparatively large and diverse sets of anatomical entities; by applying these patterns to almost the whole XAO and populating the XPO with these classes up front, our hope is to streamline curation efforts by reducing the frequency of new term requests over time.

**Table 1:**
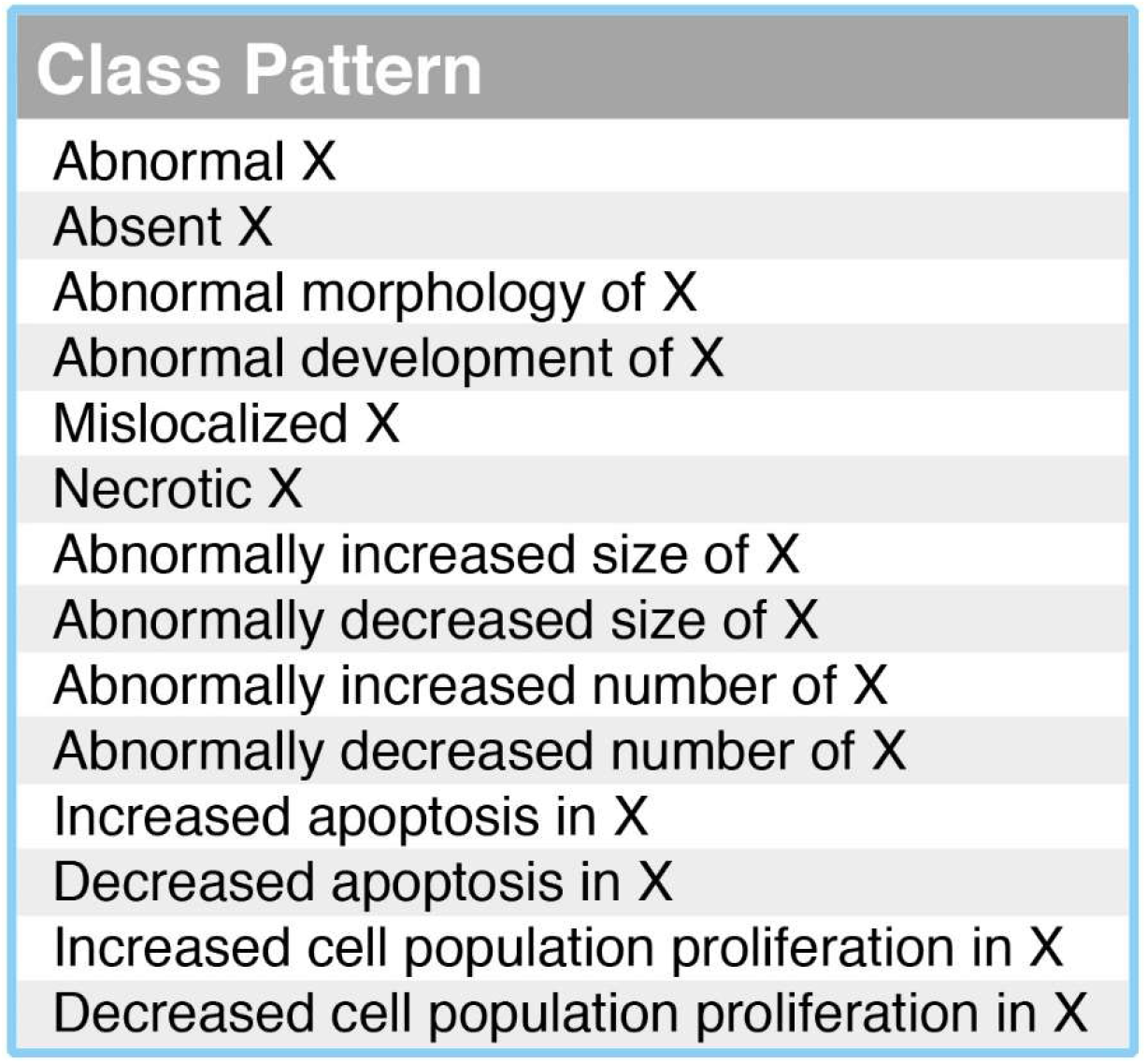
Automatically applied class design patterns for new XAO terms (X).

| Class Pattern |
| --- |
| Abnormal X |
| Absent X |
| Abnormal morphology of X |
| Abnormal development of X |
| Mislocalized X |
| Necrotic X |
| Abnormally increased size of X |
| Abnormally decreased size of X |
| Abnormally increased number of X |
| Abnormally decreased number of X |
| Increased apoptosis in X |
| Decreased apoptosis in X |
| Increased cell population proliferation in X |
| Decreased cell population proliferation in X |

In the course of developing the XPO pipeline we introduced some novel components to the ODK framework [22] approach including a system for automatically generating phenotypes from component ontologies that keeps them synchronized with the most updated patterns. If a pattern changes (e.g. a definition is updated) it is automatically updated and the ontology stays conformant. For example: the phenotype ontology reconciliation effort consortium recently decided to use the PATO class “mass density” instead of “mass”. Xenbase curators were not required to edit the XPO directly to keep it in sync with this decision, it was automatically updated by the system. In many cases in the past ontologies would employ patterns but in different ways, as illustrated by the equivalent classes for the human and mammalian (mouse) “unilateral deafness” phenotypes (Fig. 4). Even the common elements are framed distinctly with the ‘unilateral’ class being treated as a modifier only in the HPO, although it is still one of the equivalent classes in the MP. The design patterns, pipeline, and release process guarantee that ontologies developed using this system are always fully interoperable and aligned with ongoing reconciliation efforts. Consequently, the XPO and any other uPheno compliant ontologies should have identically structured equivalent classes varying only in the species-specific anatomy ontology terms used. The current XPO.v1.1 consists of approximately 22,000 terms built from a set of 80 design patterns (Supplementary Table. 2)

**Figure 4:**
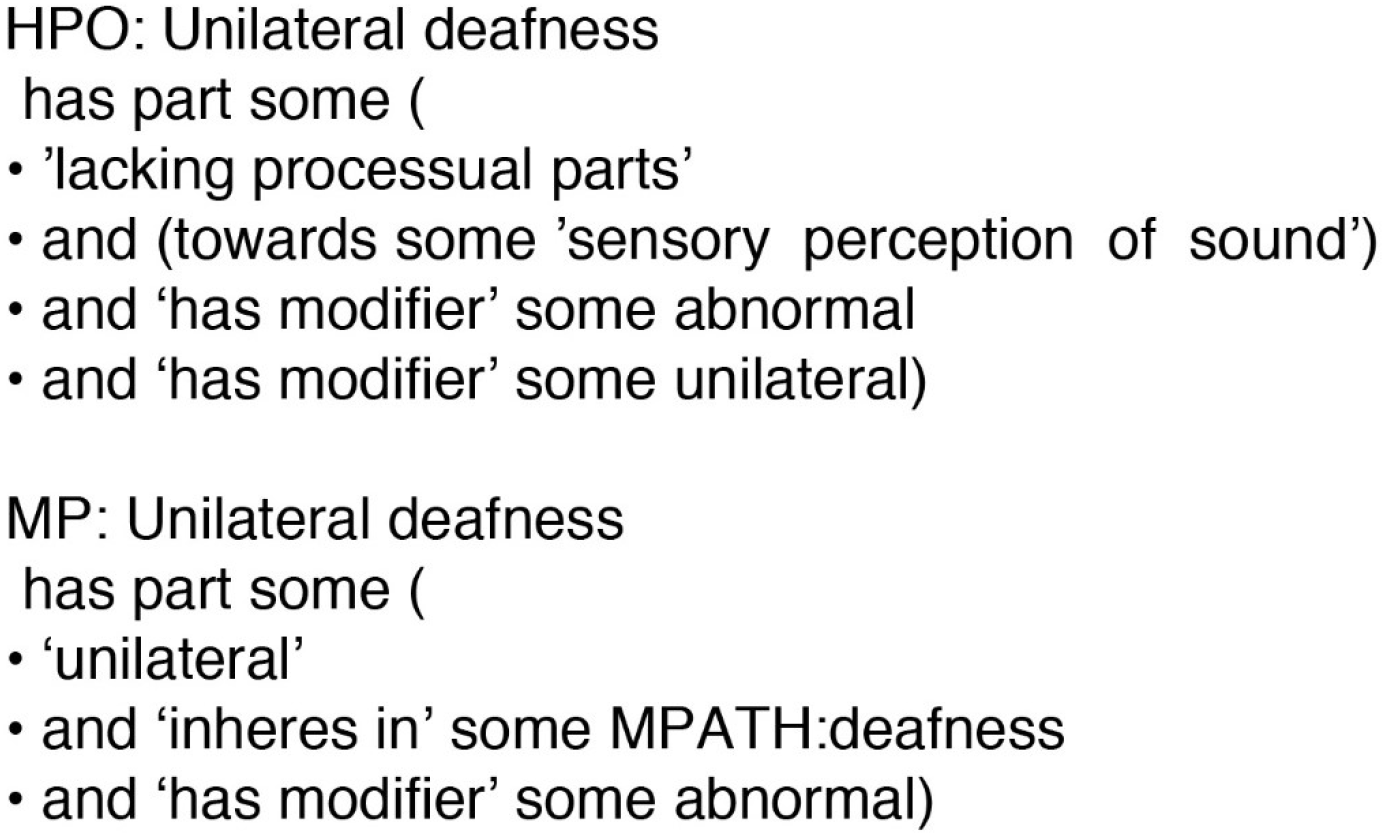
Comparison of the equivalent classes for the ‘Unilateral deafness’ phenotype class in the human (HPO) and mammalian (MP) phenotype ontologies.

### Ongoing XPO maintenance

XPO curators or community members may make requests for new XPO terms by providing the variables specified in the design pattern corresponding to the new term, in the form of IDs and labels from the relevant ontologies, as a GitHub ‘issue’. In many cases this requires only a single entity from the XAO, GO, or Neuro Behavior Ontology (NBO)[26], for instance in the “abnormalBehavior” or “edematousAnatomicalEntity” tables. More complex patterns include “in location” and “by type” components that allow the construction of phenotypes such as “abnormal axon regeneration in the optic nerve” or “Y-shaped femur in the regenerating hindlimb” without requiring the limiting and potentially time-consuming process of creating or requesting specific new classes for GO or the XAO. Additionally, TSV files are used to manage obsoleted terms and to specify which terms they have been “replaced by”, this information is also handled automatically by the pipeline to update or obsolete existing terms. New design patterns can also be requested but these are subject to the wider PORE review process and may take longer to be incorporated. There is no set release cycle for new versions of the XPO, new releases are produced when the developers consider a significant body of new terms will be generated or there is a curator need for specific terms.

### Accessing the XPO

The file structure for generating the XPO is hosted on github (https://github.com/obophenotype/xenopus-phenotype-ontology), as well as scripts and makefiles for building the XPO from source. The XPO v1.1 is available for download on Xenbase in owl (http://ftp.xenbase.org/pub/Ontologies/XPO/XPO_1.1/XPO_1.1.owl) and obo formats (http://ftp.xenbase.org/pub/Ontologies/XPO/XPO_1.1/XPO_1.1.obo) and in the Github repository. The XPO is browsable at various online endpoints such as the European Bioinformatic Institute’s Ontology Lookup Service (OLS) (https://www.ebi.ac.uk/ols/ontologies/xpo) and Ontobee (http://www.ontobee.org/ontology/xpo). The XPO is licensed under a Creative Commons CC BY 3.0 license (https://creativecommons.org/licenses/by/3.0/). The specification of the XPO in line with the ‘minimum information for the reporting of an ontology’ (MIRO) guidelines [27] is available in Supplementary Table 3.

### Application of the XPO for phenotype curation

Xenbase has begun routine curation of published *Xenopus* research articles using the XPO. The curation system allows either direct association with ‘target’ genes or indirect association through mutant or transgenic lines and reagents with existing gene associations (Fig. 5). The given example, for an image from Naert et al. 2016 [28], shows an experiment associated with two CRISPR targets captured as distinct guide RNA (gRNA) reagents. These gRNAs are associated in the database with the Xenbase genes they target, in this case rbl1 and rb1. After association of the experimental description with an image and an assay type the combined elements, stored as XB-PHENO entities, can then be associated with XPO terms such as the ‘neoplastic eye’ term for the retinoblastoma phenotype in Figure 5, or with human disease terms from the Human Disease Ontology (DO)(RRID:SCR_000476)[29]. The use of a controlled vocabulary allows the common variability in author descriptions of phenotypes to be accounted for by curator expertise so that similar or identical phenotypes are all identified with the same term. This allows the common phenotypes to be identified where a simplified text matching approach might fail.

**Fig. 5.**
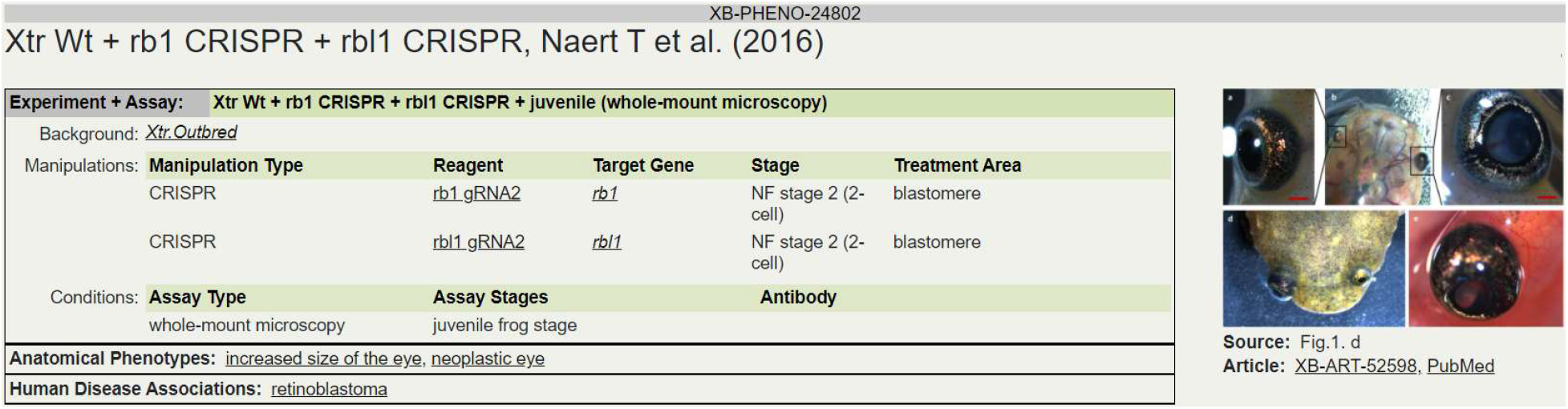
A basic phenotype curation in Xenbase. The record has a brief precis of the experiment and assay details, including background *Xenopus* strain, for the observed phenotypes as well as disease associations.

### Cross-species comparisons

Curation with the XPO allows us to link Xenopus phenotypes with and those of human, mouse, and others using the uPheno bridging ontology (Figure 6) but the linkage is currently limited by non uPheno pattern compliant terms in ontologies of other species. Mapping between non-compliant terms is not impossible, using logical or lexical mapping approaches [30], but is more challenging and often incorporates some fuzziness. Once various species phenotype data are stored in a common framework such as the Monarch SciGraph database and built using a common syntax such as the design patterns from uPheno the ease of inferring cross-species comparisons should be greatly improved.

**Figure 6:**
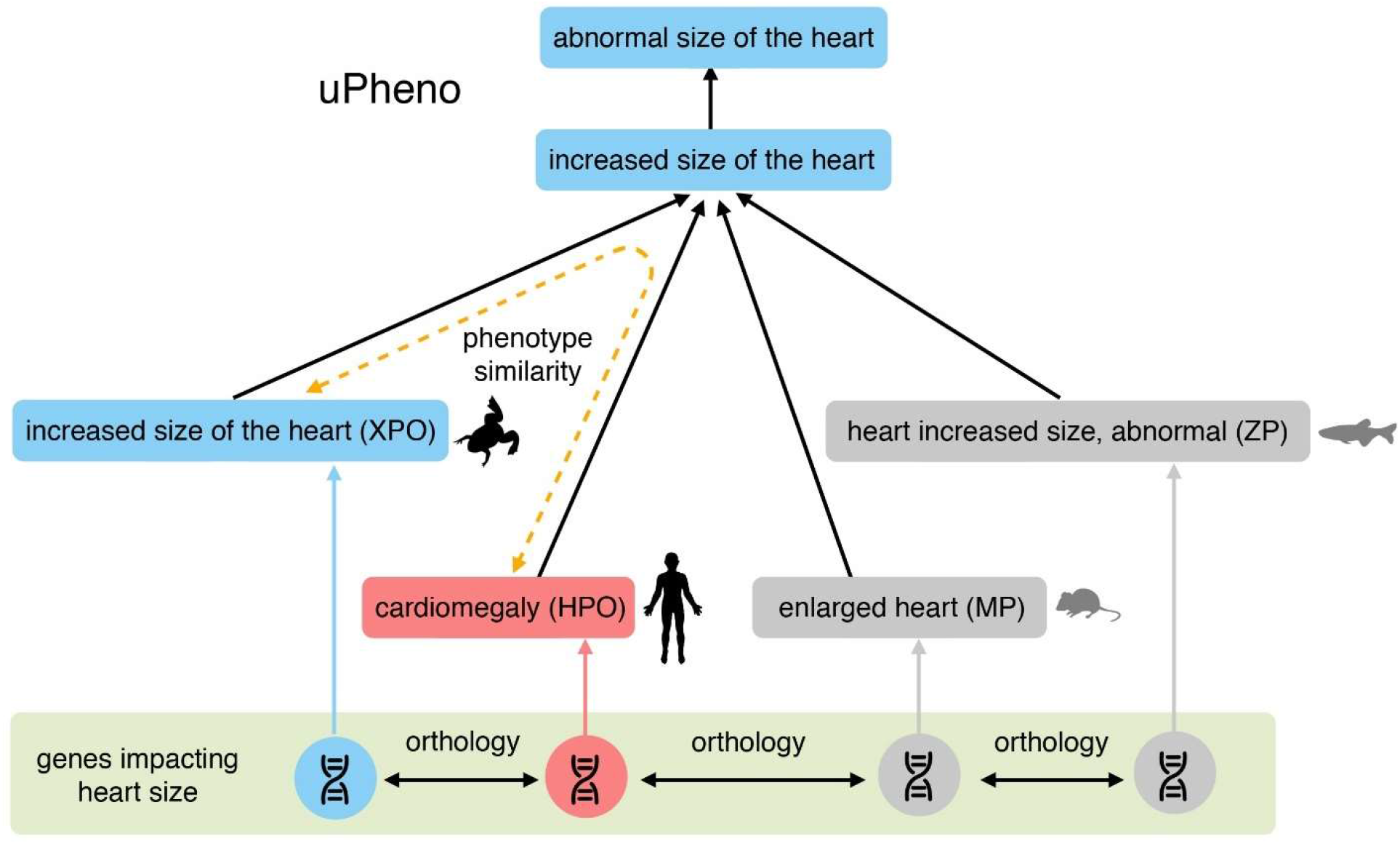
An example section of the uPheno Combined Phenotype Ontology showing the bridging term for ‘increased size of the heart’ and associated terms from 4 organism or clade specific phenotype ontologies, *Xenopus* (XPO), Zebrafish (ZP), mammalian (MP) and human (HPO). The common uPheno parent term allows the programmatic inference of phenotypic similarity (yellow dotted arrow) between the terms from differing species, and we can further infer those phenotypes caused by orthologous genes in one model organism species will give rise to similar phenotypes in humans and other species.

## Conclusions

Over the years a variety of ways of codifying certain phenotypic spectra in *Xenopus* have been put forward. These include the index of axis deficiency (IAD)[31] and the related dorso-anterior index (DAI)[32], which gave numerical values for specific degrees of axis-perturbation, and the widely used Frog Embryo Teratogenesis Assay-Xenopus (FETAX) system [33, 34], which is employed in testing the developmental toxicity of compounds and uses a standardized form to identify malformations in specific embryonic tissues and of specific types such as edema or hemorrhage. None of these existing systems provide a suitable system for general phenotype curation, either through having too narrow a focus, such as the DAI which spans a range of 0 to 10, or too shallow a capture of broader phenotypes, such as in FETAX. The new XPO provides broad coverage, incorporating several basic PATO terms for 96% of terms from the XAO, and depth down to individual cell types and subcellular components. While the XPO provides a crucial resource for internal Xenbase phenotype curation we hope it will also be a resource for researchers as a standard reference set for categorizing phenotypes, in line with this we have already produced curation for one of the broadest existing phenotypic screens in *X. tropicalis* that described and categorized phenotypes for ~136 morpholino knockdowns[35].

By building the XPO based on uPheno design patterns it is consistent with current best practices advocated by the phenotype reconciliation effort consortium to maximize interoperability. The XPO can serve as a model for ongoing efforts to integrate ontology based phenotype curation across different species and can be used as a template and workflow for the development of new phenotype ontologies [36, 37]. Refining and improving such cross-species mappings will require continued ongoing discussion between various model organism knowledgebases.

The design pattern based approach also allows the XPO to be highly responsive with new XAO terms being integrated into the XPO shortly after release. Managing new ontology requests through Github allows for transparency and anyone can submit requests allowing the research community to help direct development into areas to benefit active research. This is in line with our initial approach of basing our core terms and classes of terms for XPO development on a review and curation of existing phenotype papers to reflect the spectrum of *Xenopus* research. This research led approach reduces the likelihood of bloating the ontology as opposed to just taking every uPheno design pattern taking an anatomical term and applying it to all XAO terms and its descendants, even with basic logical restrictions on certain classes of terms as discussed previously this would still lead to rampant term proliferation.

The increased ability to perform cross-species phenotype comparisons should enhance the utility of *Xenopus* as a disease model[17]. Both the new uPheno compliant XPO and the ongoing work of projects such as Monarch to associate human diseases with Human phenotype ontology (HPO)[6] should help identifying phenotypes associated with human disease associated variants. *Xenopus* provides a rapid and flexible system for studying human sequence variants as the mRNA for a potential causative variant can be directly injected into the developing embryo [38–40] in large numbers allowing a quick survey of phenotypic effects. In addition to these forward genetics approaches, phenotypes derived from perturbing novel or under investigated genes, either by overexpression or knockdown using CRISPR or morpholinos, should be more amenable to identifying equivalent disease associated phenotypes in humans [41]. This new *Xenopus* Phenotype Ontology, along with developments throughout the biocuration community for disease and phenotype curation, will allow *Xenopus* to continue as one of the major model organisms for the study of vertebrate development and human developmental disorders and diseases.

## Supporting information

Supplemental Table 1: Design patterns proposed to PORE

Supplemental Table 2: Design patterns used in XPO

Supplemental Table 3: XPO MIRO report

## List of Abbreviations

BFO: Basic formal ontology
DAI: Dorso-anterior index
DO: Human disease ontology
EQ: Entity-Quality
FETAX: Frog Embryo Teratogenesis Assay-Xenopus
GO: Gene ontology
HPO: Human phenotype ontology
IAD: Index of axis deficiency
IRI: Internationalized Resource Identifiers
MP: Mammalian phenotype ontology
NBO: Neuro behavior ontology
OBO: Open Biomedical Ontologies
ODK: Ontology development kit
OWL W3C: Web Ontology Language
PATO: Phenotype and trait ontology
RO: Relations ontology
TSV: Tab separated values
XAO: Xenopus anatomy ontology
XPO: Xenopus phenotype ontology
ZFA: Zebrafish anatomy and development ontology
ZP: Zebrafish phenotype ontology

## Funding

This work was principally funded by grant P41 HD064556 from the Eunice Kennedy Shriver National Institute of Child Health and Human Development. Work on the uPheno template updating and revision was funded in part by the NHGRI Phenomics First Grant 1RM1HG010860-01.

## Acknowledgements

Elements of Figures 1 and 2 were adapted from Gouignard et al. 2016 [20] and used under a CC BY 3.0 license (https://creativecommons.org/licenses/by/3.0/).

Elements of Figure 5 were extracted from Naert et al. 2016 [28] and used under a CC BY 4.0 license (https://creativecommons.org/licenses/by/4.0/).

## Supplementary data

Table S1. 13 design patterns proposed to PORE

Table S2. 80 design patterns used in XPO

Table S3. MIRO report for Xenopus Phenotype Ontology

## Notes

### Competing Interest Statement

The authors have declared no competing interest.

